# The long-time orphan protist *Meringosphaera mediterranea* Lohmann, 1902 [1903] is a centrohelid heliozoan

**DOI:** 10.1101/2021.03.17.435794

**Authors:** Vasily V. Zlatogursky, Yegor Sh□shkin, Daria Drachko, Fabien Burki

## Abstract

*Meringosphaera* is an enigmatic marine protist without clear phylogenetic affiliation, but it has long been suggested to be a chrysophytes-related autotroph. Microscopy-based reports indicate that it has a worldwide distribution, but no sequence data exists so far. We obtained the first 18S rDNA sequence for *M. mediterranea* (identified using light and electron microscopy) from the West Coast of Sweden. Observations of living cells revealed granulated axopodia and up to 6 globular photosynthesizing bodies about 2 μm in diameter, the nature of which requires further investigation. The ultrastructure of barbed undulating spine scales and patternless plate scales with a central thickening is in agreement with previous reports. Molecular phylogenetic analysis placed *M. mediterranea* inside the NC5 environmental clade of Centroplasthelida (Haptista) along with additional environmental sequences, together closely related to Choanocystidae. This placement is supported by similar scales in *Meringosphaera* and Choanocystidae. We searched the Tara Oceans 18S-V9 metabarcoding dataset which revealed four OTUs with 95.5-98.5% similarity, with oceanic distribution similar to that based on morphological observations. The current taxonomic position and species composition of the genus are discussed. The planktonic lifestyle of *M. mediterranea* contradicts the view of some authors that centrohelids enter the plankton only temporarily.

---

IN 1887, Victor Hensen, a German zoologist and the founder of planktology (Lussenhop 1974) reported a finding of what he described as a yellow microscopic plant with rigid flexuous outgrowths (starr gewundenen ausläufer) in the plankton of the Baltic sea (von Hensen 1887). A few years later, a similar organism was found by Lohmann near the coast of Sicily, who erected for it the new genus *Meringosphaera* Lohmann, 1902 [1903]^1^ (from greek μ□ριγξ - bristle and σφα□ρα - ball, globe). According to Lohmann (1902 [1903] p. 68), the genus was home to unarmoured (ohne Panzer) green chromatophores-bearing cells without encircling groove (Gürtelfurche), but with long bristles for floating (Schwebborsten). In *M. mediterranea*—the species that was later fixed as type (Loeblich and Tappan 1963)— Lohmann described four chromatophores, which had a peripheral position and a cup-like shape. In contrast to the original description by Hensen, Lohmann reported green, not yellow color of the chromatophores, but he considered this species an alga of undetermined origin (protophyten unsioherer Stellung) (Lohmann 1902 [1903]). In addition to *M. mediterranea*, Lohmann included the description of three additional species. In the following years, a dozen additional species of *Meringosphaera* were described by various authors but due to a vague genus diagnosis it contained a collection of unlikely related forms, most of which were later transferred to other genera (see Silva (1979) for review of the taxonomic history).

Nevertheless, the type species—*Meringosphaera mediterranea* Lohmann, 1902 [1903]—is notable and recognizable even by light microscopy (Leadbeater 1974). It is often reported from marine plankton habitats worldwide (Table S1; Fig. 1). In some regions, it can be one of the most common and abundant planktonic species (LeRoi and Hallegraeff 2006; Thorrington-Smith 1970), reaching the concentration of 8 × 10^4^ cells l^-1^ (Booth et al. 1982). *M. mediterranea* has been reported from the surface down to 125 m deep waters (micrograph JRYSEM-305-020 on mikrotax.org) and demonstrated a temperature tolerance from 0 to 30 °C (Hallegraeff 1983; Thomsen 1982).

**Fig. 1.**
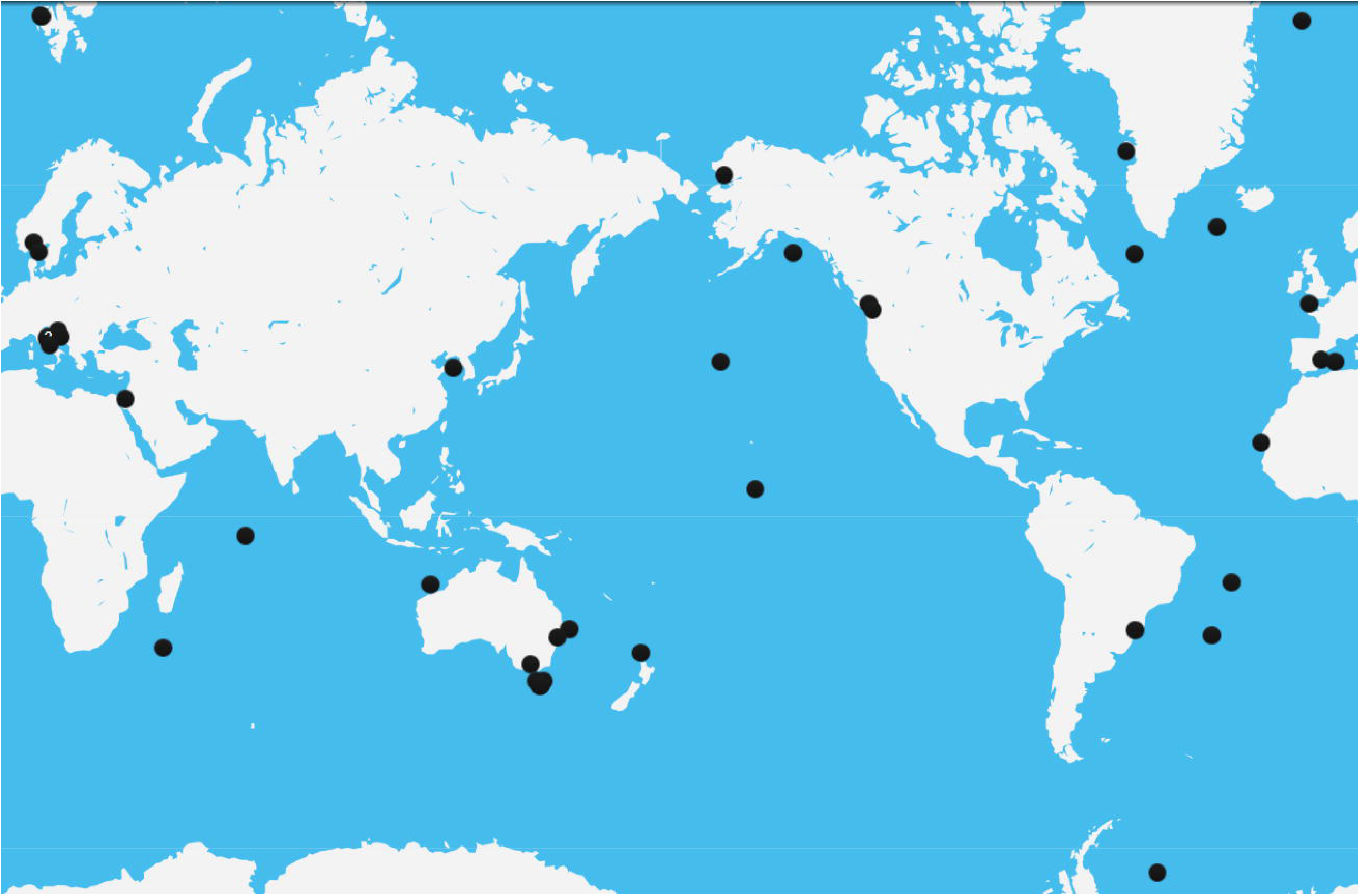
The map of the distribution of *Meringosphaera mediterranea*, based on literature records. For detailed references see Table S1.

Despite the numerous reports of *Meringosphaera* worldwide, the taxonomic affiliation of the genus has remained mysterious. Wille (1909) classified it within the green algal family Oocystaceae, based on the presence of the green chromatophores and superficial resemblance to the genera *Micractinium* and *Oocystis*. Pascher (1912; 1917; 1932) and Schiller (1916; 1925) placed the genus in the yellow-green algal order Heterococcales. This view was mostly based on the observation of two-layered siliceous walls in the cysts of *Meringosphaera triseta* (Pascher 1917), but this species was later shown to be a diatom (Throndsen and Zingone 1994). Schiller (1916) also studied *M. mediterranea* and showed that the cells are surrounded by a rigid siliceous shell (Hulle), from which siliceous bristles emerge. Each bristle arises from the low circular cup, located upward on the shell. Norris (1971) studied material from the Indian Ocean and placed *Meringosphaera* in the family Aurosphaeraceae in Chrysophyceae, based on a “golden tinge” of the living cells. Additionally, Norris provided the first ultrastructural account of the “bristles”, which were shown to be undulating tapering spine scales with short barbs directed towards the scale apex. The “shell” in turn was described as a layer of overlapping patternless plate scales with a central narrow thickening. Later, this characteristic morphology was observed and confirmed in multiple additional studies of the marine plankton. Leadbeater (1974) supported a chrysophycean affinity based on specimens collected in the Mediterranean sea, noting the similarity of the spine and plate scales with those of the chrysophyte *Chrysosphaerella* spp. Parke performed staining of the cell with dilute cresyl blue, which resulted in a rose-red color suggesting the presence of chrysolaminarin reserve products, again supporting a chrysophyte affiliation (personal communication of M. W. Parke in Leadbeater 1974). Moestrup (1979) found similar siliceous scales near the coast of New Zealand, which he also interpreted as chrysophycean affinities.

In the end of the XX century several authors expressed some doubt on the algal nature of *Meringosphaera* and suggested its possible relationship with centrohelid heliozoans. Thomsen was the first author to mention striking similarities between the scales of *Meringosphaera* with the siliceous scales of centrohelid heliozoans, particularly those of *Choanocystis perpusilla*—one of the first ultrastructurally studied (Petersen and Hansen 1960) centrohelids (personal communication of Thomsen in Moestrup (1979 pp. 65, 92). Dürrschmidt (1985) published an extensive study of the centrohelid scales and also noted that the morphology of *Meringosphaera* plate and spine scales “indicates close affinities to the heliozoa”. A similar view was expressed by Vørs (1992), who initially also suggested a relationship to the centrohelid *Choanocystis* based on the scale similarity. However, in a later report, Vørs and co-authors (1995) listed *Meringosphaera* among *incertae sedis* taxa and only vaguely referred to this organism as heliozoan-like heterotroph. The “chromatophores” were interpreted by Vørs as colored food vacuoles after preying on algae, not actual organelles. Ikävalko and Gradinger (1997) also reported a heterotrophic and centrohelid-related nature of this organism based on their observation of colorless living cells with no detectable chlorophyll, even when studied by epifluorescence microscopy with blue light excitation.

However, in most of the more recent publications, the suggestions of heliozoan-related nature were dismissed and *Meringosphaera* is referred to as chrysophyte (Fragoso 2016; Hasle and Heimdal 1998; LeRoi and Hallegraeff 2006; Liu and Chen 2015; Percopo et al. 2011; Scott and Marchant 2005; Viličić et al. 2002) or xanthophyte (Cărăuş 2002), although sometimes also as *incertae sedis* taxon (Adl et al. 2005, 2012, 2018; Bergesch et al. 2008; Bosak et al. 2012). In order to clarify the systematic position of *M. mediterranea*, we obtained the first 18S rDNA sequences for the genus based on several individual cells collected on the West Coast of Sweden, and combined these to microscopic observations of living cells and transmission electron microscopy. We unambiguously shown that *Meringosphaera* belong to centrohelids, specifically to the environmental marine clade NC5.

## MATERIAL AND METHODS

### Sampling and cell isolation

The material for this study was obtained from marine samples collected at two stations along the West Coast of Sweden: Anholt East (N 56°40’00’’ E 12°07’00’’) and Å17 (N 58°16’29’’ E 10°30’47’’). Samples were collected on 17.10.2018 at Anholt, and on 11.11.2018 and 07.12.2018 at Å17 by SMHI (Swedish Meteorological and Hydrological Institute) on the R/V Aranda, simply by collecting about 1 liter of surface water with a bucket. The water salinity at the sampling sites was around 27 psu at Anholt E and 33 psu at Å17. The water temperature was around 13 degrees in October, 10 degrees in November and around 8 degrees in December. The samples were transported to the laboratory on cooling packs in a foam plastic box. In the laboratory, the samples were passed through a 5–15 μm pore size paper membrane (VWR, Cat No. 516-0813) by gravity filtration to avoid damaging the cells– *M. mediterranea* cells are very fragile. The filters were washed in a 60 mm plastic Petri dish with 10 ml of filter-sterilized marine water. The dishes were scanned for characteristic *M. mediterranea* morphology using a 40 × lense of the Nikon Eclipse Ts2R inverted microscope, equipped with phase contrast. The detected cells were photographed with a Nikon D5300 camera and collected with a tapered Pasteur pipette. In general, we observed between 10 and 50 *Meringosphaera* cells per dish using this approach. The collected cells were placed on a glass slide for microscopy or frozen in 200 μl PCR tubes for molecular experiments. The DIC and fluorescent images were obtained from temporary preparations observed with Leica DMRXE microscope.

### Electron microscopy

Preparation of the scales for scanning electron microscopy was conducted according to Zlatogursky (2014). The cells were air-dried on the surface of a coverslip. The coverslips were washed with distilled water, attached to specimen stubs, gold-coated and observed with a Zeiss Auriga working station operated at 5 kV. Scales were measured in EM images.

### Genome amplification and PCR

Frozen single cells in PCR tubes were thawed and subjected to lysis and multiple displacement amplification (MDA) using the REPLI-g UltraFast Mini kit (Qiagen) following the manufacturer’s instructions. The product of the MDA reactions were 10X diluted and used as templates in PCR amplification of the 18S rDNA gene using broad eukaryotic primers: PF1 5′-TGCGCTACCTGGTTGATCCTGCC-3′ (Keeling 2002) and FAD4 5′-TGATCCTTCTGCAGGTTCACCTAC-3′ (Deane et al. 1998; Medlin et al. 1988). One of the obtained PCR products was purified with ExoProStar 1-Step kit (GE Healthcare US77702) and Sanger-sequenced directly at Macrogen (Netherlands). The obtained sequence was deposited in GenBank under accession number ####### (to be inserted prior to publication).

### Phylogenetic analyses

The sequence was quality-checked for ambiguous bases in Chromas Pro, and manually aligned using SeaView v. 4.3.5 (Gouy et al. 2010) on an available alignment including a broad diversity of eukaryotes. Then 1531 unambiguously aligned positions were selected for phylogenetic analysis. Initial Maximum Likelihood (ML) analyses indicated that *M. mediterranea* is a centrohelid, thus in subsequent analyses we included a broad diversity of sequences for this group. The final tree reconstruction was done using RAxML v. 8 (Stamatakis 2014), using the GTR model and 4 gamma categories to take into account across sites rate heterogeneity, after model selection in Modeltest (Posada and Crandall 1998). Assessment of clade support was performed with bootstrap resampling using 1,000 replicates.

### Search against TARA

The search against Tara oceans OTU 18S V9 v. 2 database was performed using the Ocean Barcode Atlas website (http://oba.mio.osupytheas.fr/), using our complete 18S rDNA as a query and the vsearch algorithm with 98% similarity threshold.

## RESULTS

### Light microscopy

The light microscopic description is mostly based on material collected from Anholt sampling site. The observed *M. mediterranea* cells were 4–9 μm in diameter, typically with 6–9 prominent axopodia and up to 13 undulating spine scales per optical section (Fig. 2C, D). Axopodia were distinctively granulated, 6–16 μm long, usually exceeding the cell diameter by 1.5–2 times. Cells without visible axopodia were sometimes also observed. The cells were always motionless, passively attached to the substratum or floating. All the specimens observed had a yellow-greenish tinge, containing several globular photosynthesizing bodies. One cell was squeezed with a coverslip, which revealed 6 distinct photosynthesizing bodies of about 2 μm in diameter (Fig. 2A). The chlorophyll autofluorescence emanating from the photosynthesizing bodies was clearly visible when subjected to fluorescence microscopy with blue excitation and green-red emission (Fig. 2B).

**Fig. 2.**
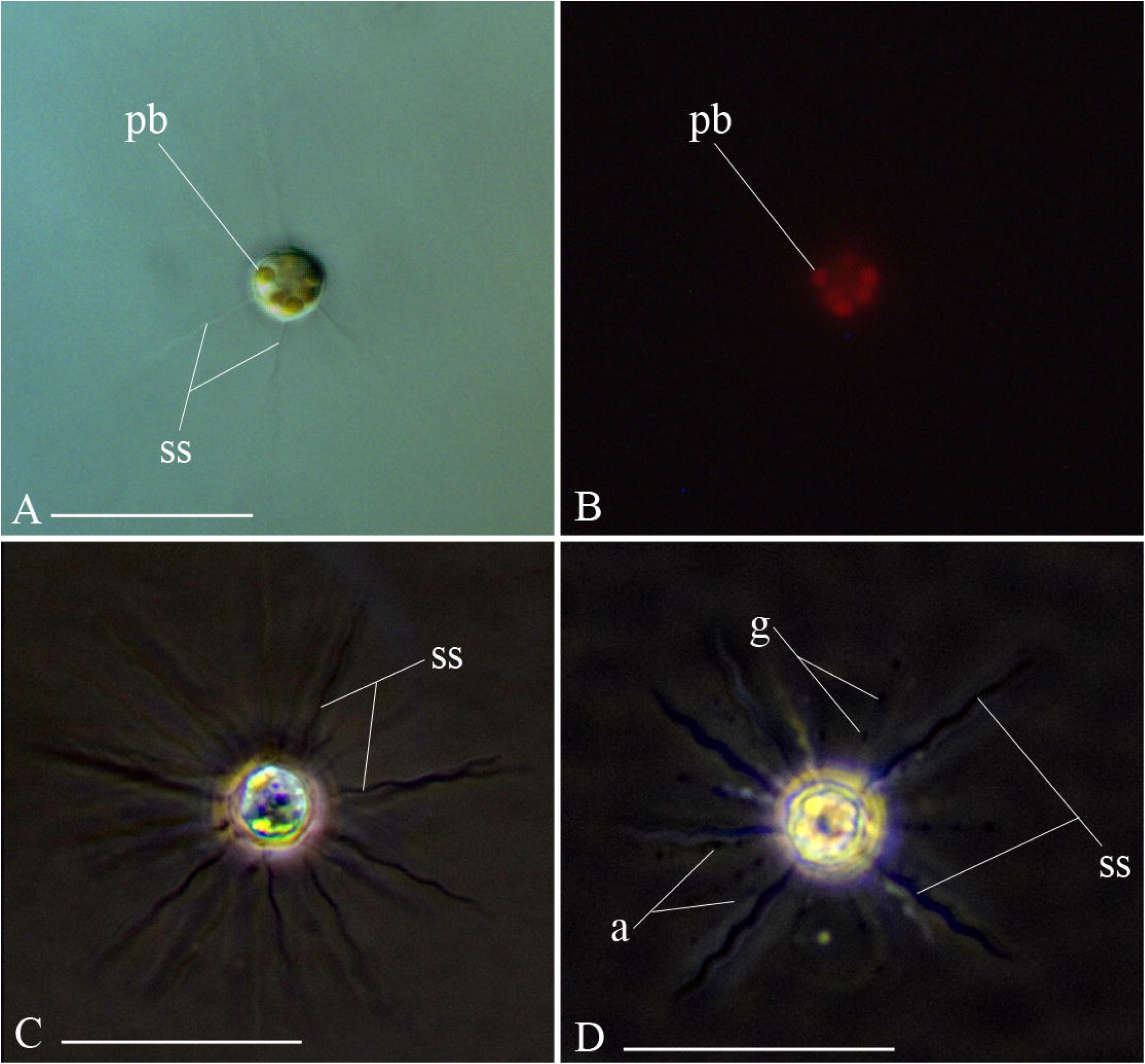
*Meringosphaera mediterranea*, collected in Å17 (**A, B**) and Anholt (**C, D**) sampling points. Light microscopy, general view of the living cell. **A**. Differential interference contrast (DIC). **B**. The same field as in (**A**), fluorescent microscopy, blue excitation and green-red emission filters. **C**. Phase contrast, the cell is squashed under a cover-slip. **D**. Phase contrast. Undisturbed cell in a Petri dish. Abbreviations: a - axopodia; g - granules; pb - photosynthesizing bodies; ss - spine scales. Scale bars 20 μm.

### Electron microscopy

The skeletal elements of the cells (plate scales and spine scales) from Å17 (mostly) as well as from Anholt sampling points were studied with scanning electron microscopy (Fig. 3). The spine scales were typically 16–25 μm long, but in one case a single 31 μm giant spine scale was observed. The shaft of each scale with 9–13 undulations was covered with barbs (Fig. 3A, B). Usually, there were 1–3 longer barbs, about twice longer than the shaft diameter at the proximal part of the scale (Fig. 3C) and multiple (15–24) shorter barbs distributed in a helicoidal pattern along the whole scale length. Shorter barbs were flattened, triangular, and slightly curved in the direction of the scale tip. Spine scale shafts were hollow inside with internal septa, which seemed to be located at the inflection points between undulations. The bases of the spine scales were convex, with a single indentation and multiple (10–13) short teeth along the margin. The plate scales were patternless, oval 1.8–3.2 × 1.4–2.1 μm, with a short central thickening.

**Fig. 3.**
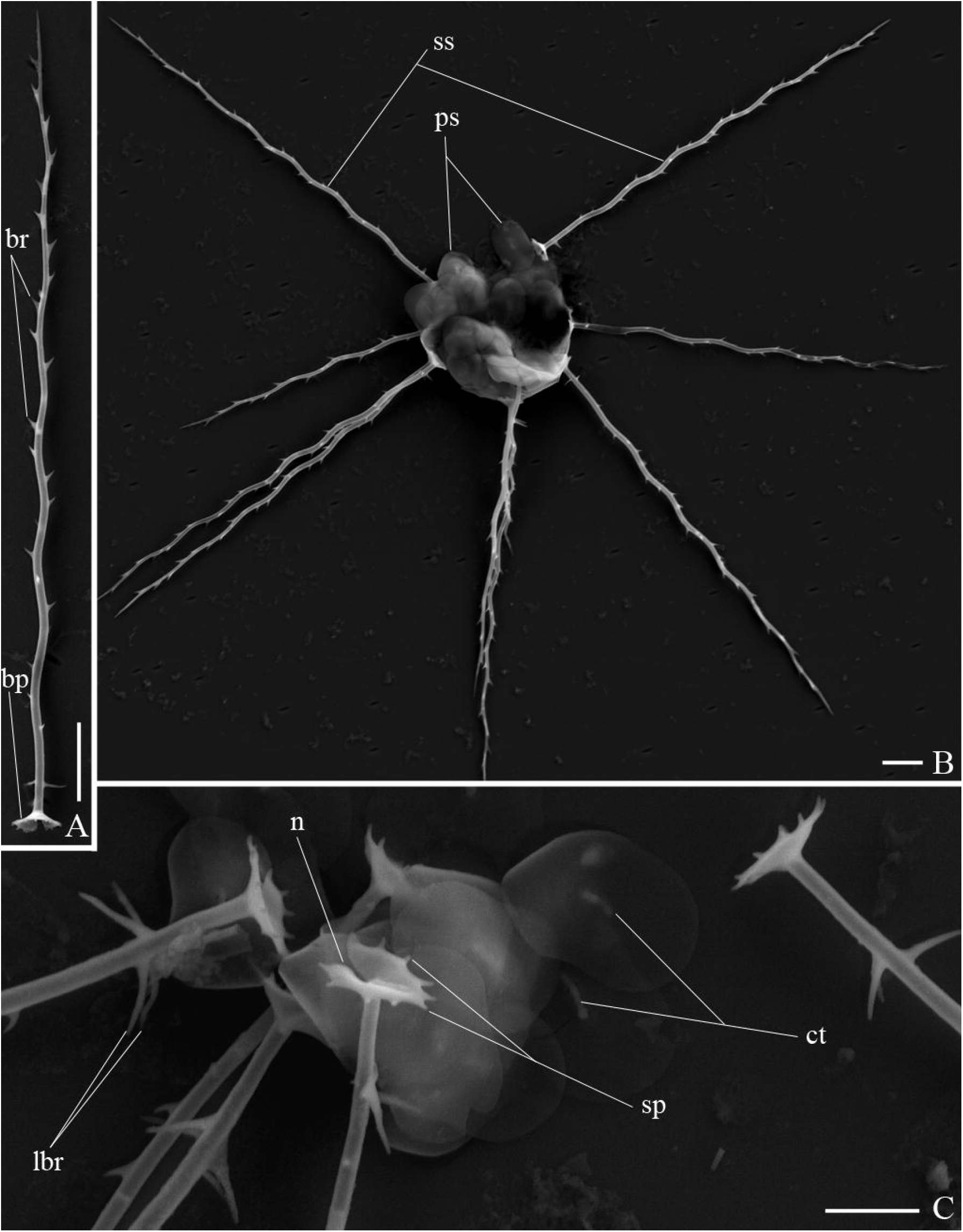
*Meringosphaera mediterranea*, collected in Å17 sampling point. Scanning electron microscopy. **A**. Individual spine scale. **B**. General view. **C**. Close up of plate scales and spine-scales proximal parts. Abbreviations: bp - basal plate of the spine scale; br - barbs; ct - central thickening of the plate scale; lbr - elongated barbs on the proximal part of the spine scale; n - notch on the spine scale base; ps - plate scales; sp - spikes on the spine scale base; ss - spine scales. Scale bars: **A, B** - 2 μm; **C** - 1 μm.

### Molecular phylogeny

A good quality sequence of the 18S rDNA gene of 1720 bp was obtained from SAG SC462 from Anholt, which we considered the first sequence for the genus *Meringosphaera* (see discussion). A comparison in GenBank by BLASTn for highly similar sequences returned only unnamed environmental sequences belonging to centrohelids. The best environmental hit was KF130174 with 99.65% similarity and the best named hit - *Chlamydaster sterni* KY857824 with 93.62% similarity. In order to place this sequence in a phylogenetic context, a tree reconstruction including a broad selection of centrohelid taxa was performed. This analysis recovered a moderately supported position (81% bootstrap support) of the sequence within the NC5 environmental clade of Pterocystida (Fig. 4), following the environmental clade nomenclature of Cavalier-Smith and Chao (2012) and Sh□shkin et al. (2018). The NC5 clade included several additional environmental sequences (of a marine plankton origin wherever specified) and was sister to the Ch1 clade, containing the closest group with named species Choanocystidae.

**Fig. 4.**
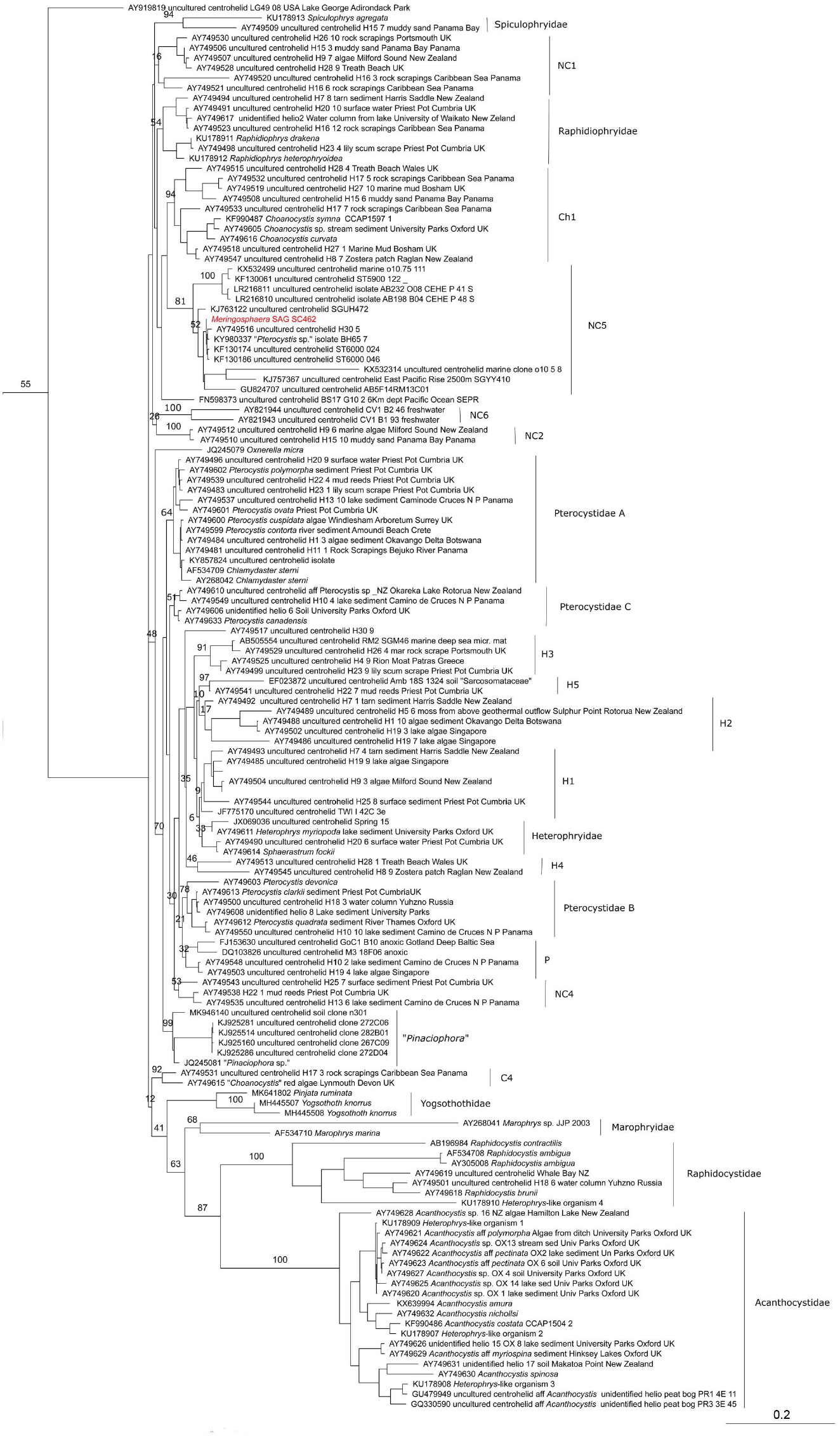
Maximum likelihood tree for 18S rDNA of 141 centrohelids and outgroup of 57 sequences (1531 sites; GTR; bootstrap 1,000 replic. 4 rate classes). Outgroup and support values for shallow clades have been removed for clarity.

### Metabarcoding dataset search

We investigated the oceanic distribution of the *Meringosphaera* sequence derived from SAG SC462. A search against the Tara Oceans 18S V9 v. 2 metabarcoding dataset returned four hits (OTUs 18Sv.9-v.2 346365; 18Sv.9-v.2 557845; 18Sv.9-v.2 1004; 18Sv.9-v.2 35493), which had a global distribution (Fig. 5), maximal relative abundances in surface waters, but were also quite common in the vicinity of the deep chlorophyll maximum. In general, these OTUs had a low relative abundance, representing at most 2.33e^-3^. In the mesopelagic zone, the OTUs were less abundant but still present worldwide. The temperature tolerance range was high, varying from 2 to 33 □, while salinity tolerance was quite narrow, varying between 32 and 41 ppt and never below 28 ppt (Fig. 5). This matched our own observation that *Meringosphaera* occurred frequently on the Swedish West coast where the salinity is at about 33 ‰, but could never be detected in samples from the Baltic Sea proper with a much lower salinity. The OTUs were found in all the size fractions, except the finest (0.8–3 μm).

**Fig. 5.**
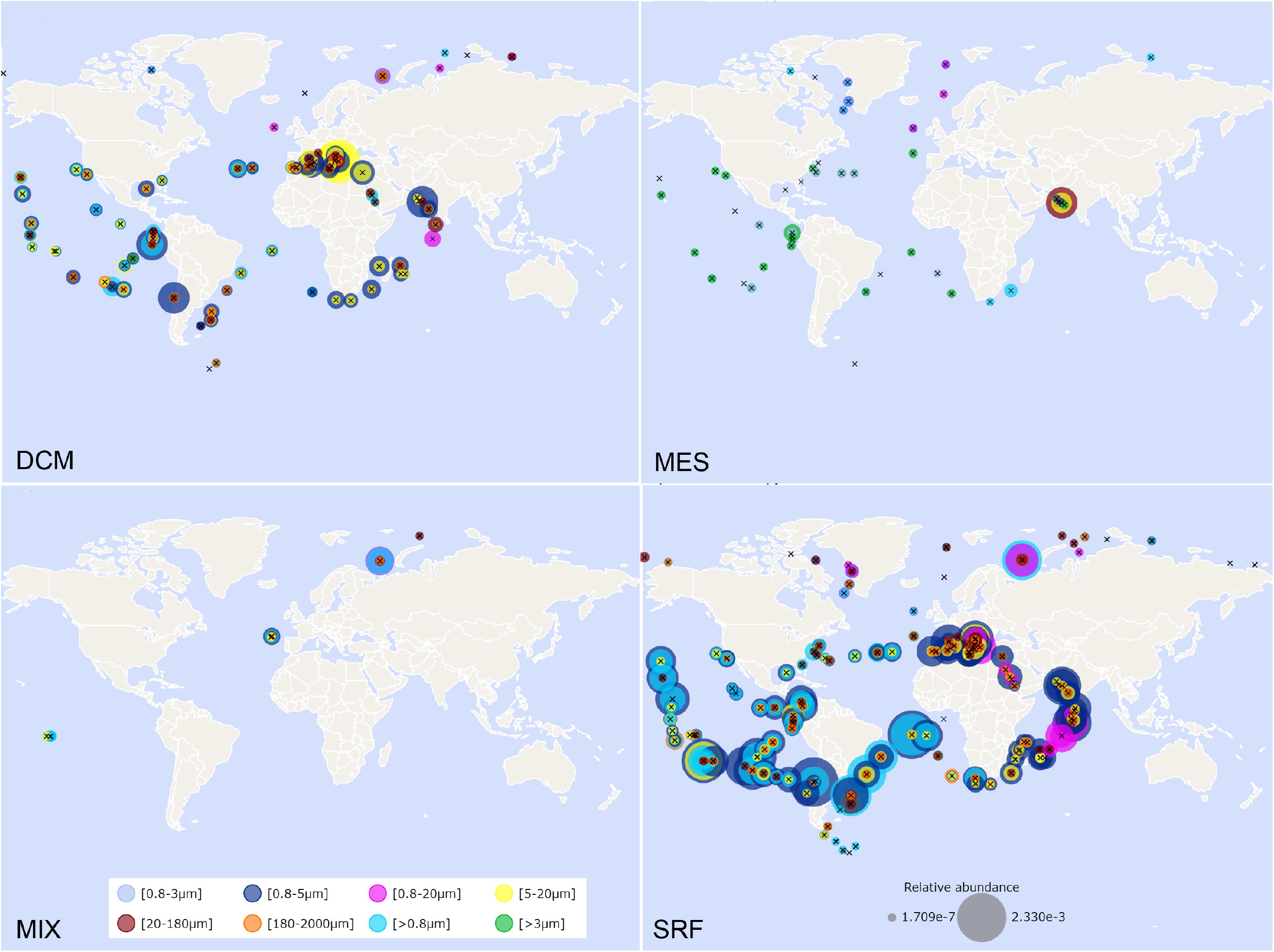
Map of the geographic distribution of four *Meringosphaera-*related OTUs (Tara Oceans 18S V9 v. 2). Circles are proportional to abundance. Plankton organismal size fractions are color-coded. DCM - deep chlorophyll maximum; MES - mesopelagic zone (200-1000 m); MIX - mixed layer; SRF - surface water.

## DISCUSSION

The morphological analysis by light microscopy and the ultrastructure of the specimens studied strongly suggests that these organisms correspond to *Meringosphaera mediterranea*. Most of the species described under the genus name *Meringosphaera* have eventually found their homes in other genera, considered synonymous or too distinct and/or poorly described to safely keep them in the genus (Table 1). In fact, Silva (1979) proposed that the species diversity of the genus *Meringosphaera* should be restricted to *M. mediterranea* and *M. aculeata* Pascher, 1932. *M. aculeata*, despite the considerable similarity to *M. mediterranea*, is distinct in having fewer undulations per scale and very long barbs, which Wulff (1919) was able to detect with light microscopy. His fig. 14 taf. II is in a good agreement with type 2 scales on scanning electron micrographs (fig. 4 and 5 of Norris (1971)), as well as with the micrograph JRYSEM-317-330 published on the microtax.org website. Other micrographs, published under the name *M. mediterranea* are quite heterogeneous and probably represent several closely related species. For example, the morphotype with stellate base of the spine scale is very distinctive (fig. 3 and 8 of Norris (1971); fig. 2 of Vørs and co-authors (1995); micrograph JRYSEM-260-16 on microtax.org). The cells in our study were similar to typical *M. mediterranea* as in fig. 1 of Norris (1971); Plate 4 B–F of Leadbeater (1974); fig. 6 of Vørs (1992) and many other published micrographs, both by details of the ultrastructure and morphometric characters.

**Table 1.** The list of all the species described in the genus *Meringosphaera* and their final taxonomic homes.

| Species names (chronologically) | Final destiny |
| --- | --- |
| ***M. mediterranea* Lohmann, 1902[1903]** | type, valid |
| *M. baltica* Lohmann, 1902[1903] | Synonym of *M. mediterranea* acc. to (Lohmann, 1908) |
| *M. divergens* Lohmann, 1902[1903] | Transferred to *Sciadosphaera* by (Pascher, 1938) |
| *M. hydroidea* Lohmann, 1902[1903] | Transferred to *Ophiaster* by (Lohmann, 1913) |
| *M. serrata* Lohmann, 1908 | Presumable coccolithophorid acc. to (Pascher, 1932) |
| *M. radians* Lohmann, 1908 | Transferred to *Apedinella* (Pedinellida) by (Campbell, 1973) |
| *M. henseni* Schiller, 1916 | Presumable separate genus acc. to (Silva, 1979) |
| *M. trisetata* Schiller, 1916 | Synonym of *Chaetoceros trondsenii* var. *trisetosa* (Bacillariophyceae) acc. to (Thronsen and Zingone, 1994) |
| *M. merzii* Schiller, 1925 | Presumable separate genus acc. to (Silva, 1979) |
| *M. tenerrima* Schiller, 1925 | Transferred to *Raphidosphaera* by (Silva, 1979) |
| *M. setifera* Schiller, 1925 | Transferred to *Raphidosphaera* by (Silva, 1979) |
| ***M. aculeata* Pascher, 1932** | valid |
| *M. brevispina* Pascher, 1932 | Transferred to *Raphidosphaera* by (Silva, 1979) |
| *M. sol* Pascher, 1932 | Transferred to *Actinellipsoidion* (Xanthophyceae) by (Ettl, 1977) |
| *M. wulffiana* Pascher, 1938 | Transferred to *Raphidosphaera* by (Silva, 1979) |
| *M. spinosa* Prescott, 1949 | Not a member of *Meringosphaera* complex acc. to (Silva, 1979) |

**Fig. 6.**
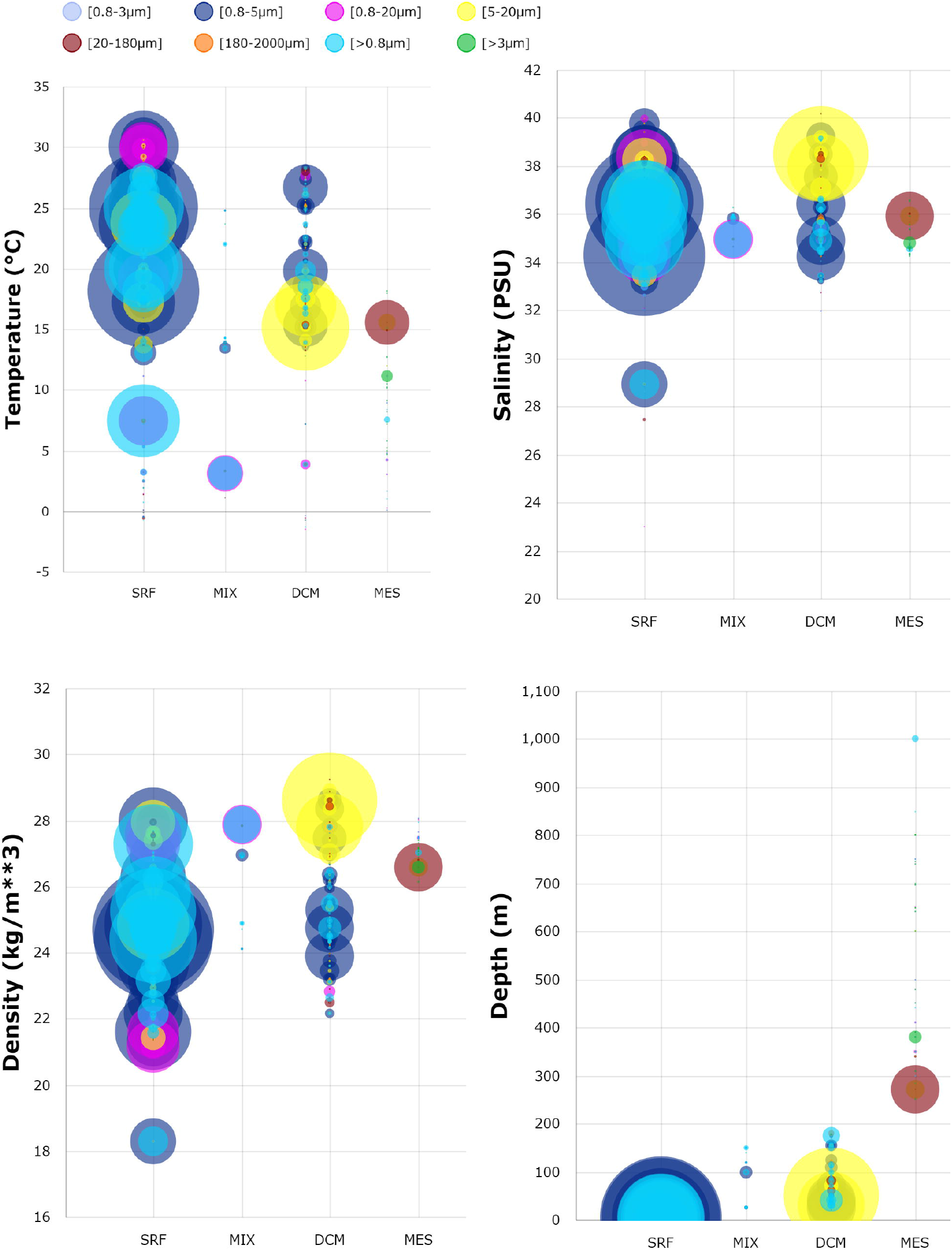
Bubble plots, representing the co-variation of four *Meringosphaera-*related OTUs (Tara Oceans 18S V9 v. 2) abundance and an environmental feature at four sampling depth fractions (DCM - deep chlorophyll maximum; MES - mesopelagic zone (200-1000 m); MIX - mixed layer; SRF - surface water). Circles are proportional to abundance. Plankton organismal size fractions are color-coded.

The new *M. mediterranea* sequence was clearly positioned in the environmental NC5 group in centrohelids. In total, the NC5 group now contains 12 environmental sequences in GenBank, all marine, in addition to the new *Meringosphaera* 18S rDNA sequence. Our phylogeny also suggested a weak relationship to Ch1 - a mostly environmental clade with only two morphologically characterized sequences, representing the family Choanocystidae. Although this relationship requires confirmation due to the low bootstrap support, it is in agreement with the view of some authors (Moestrup 1979; Vørs 1992) who emphasized the similarity in the scale structure between *M. mediterranea* and *Choanocystis* spp. Since *Choanocystis* is a very species-rich genus with 18 described species (Mikrjukov 1995; Tihonenkov and Mylnikov 2010; Zlatogursky 2010, 2014), only two of which have been sequenced (Sh□shkin et al. 2018), it is possible that some of the morphotypes attributed to *Choanocystis* actually belong to the NC5 clade. This is even more probable for the four exclusively marine species. For example, *Choanocystis antarctica* Tihonenkov et Mylnikov, 2010 is one of the best candidates to be closely related to *Meringosphaera* since it also has a typical barb on each spine scale and each spine scale in this species possesses a single undulation.

The search using the new *M. mediterranea* sequence against the Tara Oceans 18S V9 v. 2 database returned four OTUs with > 98% similarity, which we attribute to the same species. The geographic distribution of these OTUs is in good agreement with the morphology-based reports of *M. mediterranea*, confirming that this species is a global member of the oceanic plankton communities (compare Fig. 1 and Fig. 5). The finding of a global planktonic centrohelid species is at odds with the idea that centrohelids are only temporarily found in the water column. Mikrjukov (2002) argued that centrohelids are permanent important consumers in both freshwater and marine benthic communities but are only temporarily playing key ecological roles in the plankton, usually twice a year for a month. The common occurrence of *M. mediterranea* in many localities instead indicates that some centrohelids at least represent permanent members of planktonic communities. Our Tara Oceans search also confirmed a broad temperature tolerance as noted by Hallegraeff (1983), but a salinity tolerance restricted to oceanic values (Fig. 6). Finally, the vertical profile of *Meringosphaera-*related OTUs showed a distribution from the surface to mesopelagic zone, with relative abundance up to 2.27e^-3^ at deep chlorophyll maximum, which may be correlated to the presence of photosynthetic bodies in this species (Fig. 5, 6). The nature of these photosynthetic bodies is one of the most outstanding questions regarding *Meringosphaera*, but unfortunately we failed to obtain sequence data that could help identify their origin. These bodies could correspond to transient associations such as kleptoplasts or facultative symbionts, stable endosymbionts, or even permanent photosynthetic organelles. Regardless of the final answer on these bodies, here we’ve undoubtedly placed an important player in the marine planktonic ecosystem to its correct phylogenetic home.

## ACKNOWLEDGEMENTS

This study was supported by a scholarship from Carl Tryggers Stiftelse to VZ (PI: FB) and the Russian Science Foundation grant no. 20-74-10068. Equipment of the core facility centers “Resource Center for Nanotechnology”, “Development of molecular and cell technologies”, “Computing center SPbU” and “Culturing of microorganisms” of Saint Petersburg State University was applied. FB was supported by a grant from Science for Life Laboratory. Authors are grateful to Ann-Turi Skjevik for providing SMHI samples.

## SUPPORTING INFORMATION

**Table S1**. The summary of literature reports of *Meringosphaera mediterranea*.

## Footnotes

1 The volume with his work was issued with the year ‘1903’ on the title page, but according to bibliographic notes of Oltmanns (1903, column 210), Zschokke (1903, p. 324), Matzdorff (1903a p. 116; 1903b p. 191), Graf zu Solms Laubach and Oltmanns (1904 column 103), and Krumbach (1907 p. 463), the actual year of its issue is 1902.

